# The Effects of Environmental Enrichment on the Behavior of Laboratory Zebrafish (*Danio rerio*)

**DOI:** 10.64898/2026.10.28.762569

**Authors:** Beatrice C. Rejouis, Kristen E. Severi

## Abstract

Enclosures for laboratory animals are often simple for economic and practical reasons. Enrichment items, such as plants, toys, and refuges, have been shown to improve measures of welfare in commonly used species such as mice and dogs. The numbers of laboratory zebrafish have been increasing in recent years, and most are housed in systems devoid of enrichment. The aim of this study is to assess the impact of environmental enrichment on adult laboratory zebrafish behavior. Experimental fish were raised in the presence of three different enrichment regimes: plastic plants, toys, and photos of gravel. These were compared to a control group raised under standard housing conditions. Once the fish reached sexual maturity, we measured aggression, reproduction, and growth. We found that each enrichment treatment led to significant reductions in aggression in a mirror test compared to control. Reproduction was largely unaffected by the three enrichment treatments. However, we found egg production was decreased following the gravel treatment, and egg survival was increased following the plant treatment. There were no significant impacts of enrichment on measures of growth. In a parallel experiment, breeding fish were exposed to plastic plant enrichment during overnight breeding. Surprisingly, there was no significant impact on egg production or egg survival.

## Introduction

Laboratory animals typically live in less complex environments than they experience in their natural habitat. Specifically, laboratory environments reduce the mobility of the animal and complexity of environmental stimuli. These differences in habitat have been shown to have negative impacts on the behavior, health, and welfare of the animals (Geret et al., 2011; Hansen & Berthelsen 2000; Williams et al. 2009). The addition of simple enrichment items, however, has been shown to have significant positive impacts on animal welfare in laboratory settings (Wafer et al., 2016; Woodward et al., 2019; Williams et al. 2009). Zebrafish are growing as an animal model with the default housing consisting of a bare tank for ease of identification and monitoring (Kinth et al. 2013; Williams et al. 2009). This paper assesses the impact of simple enrichment regimes that are compatible with standard housing in a laboratory setting.

Zebrafish are a popular animal to study in captivity for numerous reasons. Their behavior has been well defined with distinct actions that can be attributed to events such as aggression or courtship (Leshner and Pfaff, 2011). In addition, zebrafish offspring begin life as transparent embryos, making observations of their inner workings simple and noninvasive (White, 2008). They live in communal groups where a typical housing unit would hold about 5 fish per liter of water, with each individual growing to be about 2-3 centimeters in length (Singleman and Holtzman, 2014). These factors, along with their rapid spawn rate, make zebrafish commonplace in research laboratories.

This study aims to measure the impact of long-term exposure to environmental enrichment items during growth on adult zebrafish. Enrichment items are objects intended to encourage normal species behavior or otherwise stimulate the subject (Woodward et al., 2019). Some animals in captivity can benefit from these additions, however, it should not be assumed that this benefit is universal across species. Some studies suggest enrichment could increase fecundity, or the number of eggs a pair of breeding zebrafish produce (Wafer et al., 2016; Woodward et al., 2019). Yet, other studies suggest enrichment items increase aggression in male adult zebrafish (Lee et al., 2018; Schroeder et al., 2014; Woodward et al., 2019). In-tank aggression would endanger the research animals and should be avoided at all costs. Yet a greater overall yield when breeding would save labor and could reduce the number of adult animals requiring housing. We hypothesized that long-term environmental enrichment would increase aggression in adult zebrafish, as well as increasing zebrafish growth and breeding effectiveness, relative to zebrafish raised without enriched tanks. We specifically expected to observe an increase in the number of eggs laid, as was demonstrated in previous experiments on the topic (Wafer et al., 2016; Woodward et al., 2019). However, we found in this study that long-term environmental enrichment decreased aggression in adult zebrafish toward their conspecifics and that only certain types of enrichment increased zebrafish breeding effectiveness.

### Ethical Review

The work described in this manuscript was reviewed and approved by Rutgers University-Newark IACUC. The associated protocol was titled “Environmental enrichment in zebrafish habitats”.

## Materials and Methods

### Study Design

In this experiment, adult zebrafish raised in the presence of different forms of environmental enrichment were compared to each other and a control group raised without enrichment. One group was raised in the presence of plastic plants, another in the presence of plastic toys, another with images of gravel and a final group without any treatment, the control. Each group had 2 tanks of about 25 larvae each at their disposal. At 100 dpf, the fish in each tank were measured and sexed. At 160 dpf, 26 fish were selected from each tank to be cross bred for the reproduction tests. These reproduction, or breeding tests, were repeated after 244 dpf. After 245 dpf the fish were measured and sexed again. At 18 months 6 fish were randomly selected from each treatment and their aggression was measured using a mirror test. After the conclusion of the aggression tests, the data was analyzed.

### Sample Size

About 50 larvae were used in each sample group for a total of about 200 larvae used. 151 juveniles survived until the measurements at 100 dpf (34 fish in the control group, 36 in the plant group, 39 in the toy group and 42 in the gravel group). 104 adults from this same pool were used in the breeding tests with 26 fish chosen from each group. This same group of adults were then measured at the conclusion of the breeding tests. 6 adult fish were then randomly selected from the pool for the aggression tests.

The numerical results from each trial were compiled into a spread sheet and then analyzed with estimation statistics (Ho et al., 2019, estimationstats.com). A p value of less than 0.05 was considered statistically significant.

### Inclusion and Exclusion Criteria

‘All fish involved in the study which survived until the point of the experimental tests were included in the experiment. No specific exclusion or inclusion criteria was established prior. In some cases during the reproduction tests, the pairs failed to breed, however the data was still noted. During measurement and sexing after 245 dpf some of the fish could not be identified as either male or female with a high degree of certainty. For this reason corrected values are included in the tables reporting the results from the reproduction tests only using data from the pairs that could be visually confirmed to contain one male and one female.

### Randomization

Fish were randomly assigned to a treatment group at 0 dpf. The eggs were separated into their groups and maintained in those groups until the conclusion of the study. If a procedure was to be done fish from the control group would be handled first, followed by the plant group, then the toy group and finally the gravel group. Measurements and sexing was done tank by tank all in the same day within the same period of time. Breeding tests proceeded with 2 pairs from each group crossed overnight at the same time. The final breeding test proceeded with 3 pairs from each group. The breeding tanks were placed in a similar orientation each time and egg collection and return of fish to their home tank proceeded in the same procedure order detailed earlier, starting with the control group and ending with the gravel group.

The fish were kept in the same general location with the only changes in location being those related to the experiment, such as a fish being moved to a crossing tank for breeding or anesthetization for measurement. The fish would occasionally be moved from tank A of their experimental group to tank B to denote completion of a particular test, but there would always be a minimum of 4 fish in each tank. Any additional confounders were not specifically controlled.

### Blinding

All researchers and animal care takers were aware of what fish belonged to which group at all times during the experiment including data collection and interpretation. The experimental groups were labeled on each tank and crossing tank when the fish were removed from the colony for breeding.

### Outcome Measures

Growth and breeding tests utilized the maximum number of participants available. Both were assessed objectively based on the length of the fish or the number of eggs obtained and the number that subsequently survived. Aggression tests used a small pool and correlated location to the level of aggression.

### Statistical Methods

The numerical results from each trial were compiled into a spread sheet and then analyzed with estimation statistics (Ho et al., 2019, estimationstats.com). This approach uses bootstrapping to determine 95% confidence intervals and show how the results would be represented in a larger sample size.

### Experimental Animals

The zebrafish used in these experiments were housed at the New Jersey Institute of Technology (NJIT) on a recirculating system (ActiveBlue, Tecniplast). The room was set to a 14-hour light:10-hour dark light cycle and the system was maintained at 28.0 °C.

All fish were of the AB WT strain, spawned from a group cross of AB WT siblings. We included a mixture of siblings and non-siblings in each tank. From 0-5 dpf eggs, embryos, and larvae were housed in petri dishes of fish facility water in a 28.5 °C incubator with a 12:12 light:dark cycle. During this time they were not fed and one third of the water per dish was changed daily. From 5 to 35 dpf they were housed in a bare 3.5 liter tank with a net insert (Tecniplast, USA) and fed both powder and live rotifer food. Powder food was Gemma Micro (Skretting, USA) and the size of the Gemma pellets and number of portions fed per day were chosen according to the guidelines suggested by Skretting through their own research (Lawrence et al., 2012). Live rotifer food consisted of approximately 1.5 ml of approximately 300-500 live rotifers per milliliter of water (Reed Mariculture, *Brachionus spp.*).

From 5 to 35 dpf they were given about 0.01 grams of Gemma Micro 75 powder (Skretting, USA) for three daily feeds, each at least three hours apart. Additionally, rotifers were fed following each powder feed. From 35 dpf to 90 dpf, the fish were fed 0.01 grams of Gemma Micro 300 powder twice a day and at least one feeding of rotifers per day. After 90 dpf, when the fish were adults, they were fed 0.01 grams of Gemma Micro 500 pellets twice a day and only fed rotifers once daily. Their feeding regimen was recorded throughout the experiment.

### Experimental Procedure

#### Enrichment

The experiment consisted of four groups of fish: one group as a control and three groups which were introduced to environmental enrichment. Each group contained two tanks of fish, with 20-25 individuals in each tank. The fish were all raised in the same manner, with the only difference between them being the enrichment contents of the tank.

At 35 dpf, enrichment items were introduced to the six experimental tanks (**Figure 1**), and all tanks were visually separated from each other with an opaque laminated screen in between the tanks. Environmental enrichment in this study consisted of artificial plants, rubber toys, and images of gravel (**Figure 1**). We substituted live plants with artificial plants to reduce the project’s impact on other fish housed on the system. Artificial plants have been shown to be effective enrichment tools and they reduce the variability caused by using live plants (Collymore et al., 2015; Lee et al., 2018; Wafer et al., 2016). The rubber toys simulated the caves and hideaway areas often provided in home aquariums. Lastly, images of gravel have been proven as effective enrichment tool and were used in place of aquarium gravel to reduce the chance of injury to the fish and or buildup of debris in the tank environment (Lee et al., 2018).

**Figure 1.**
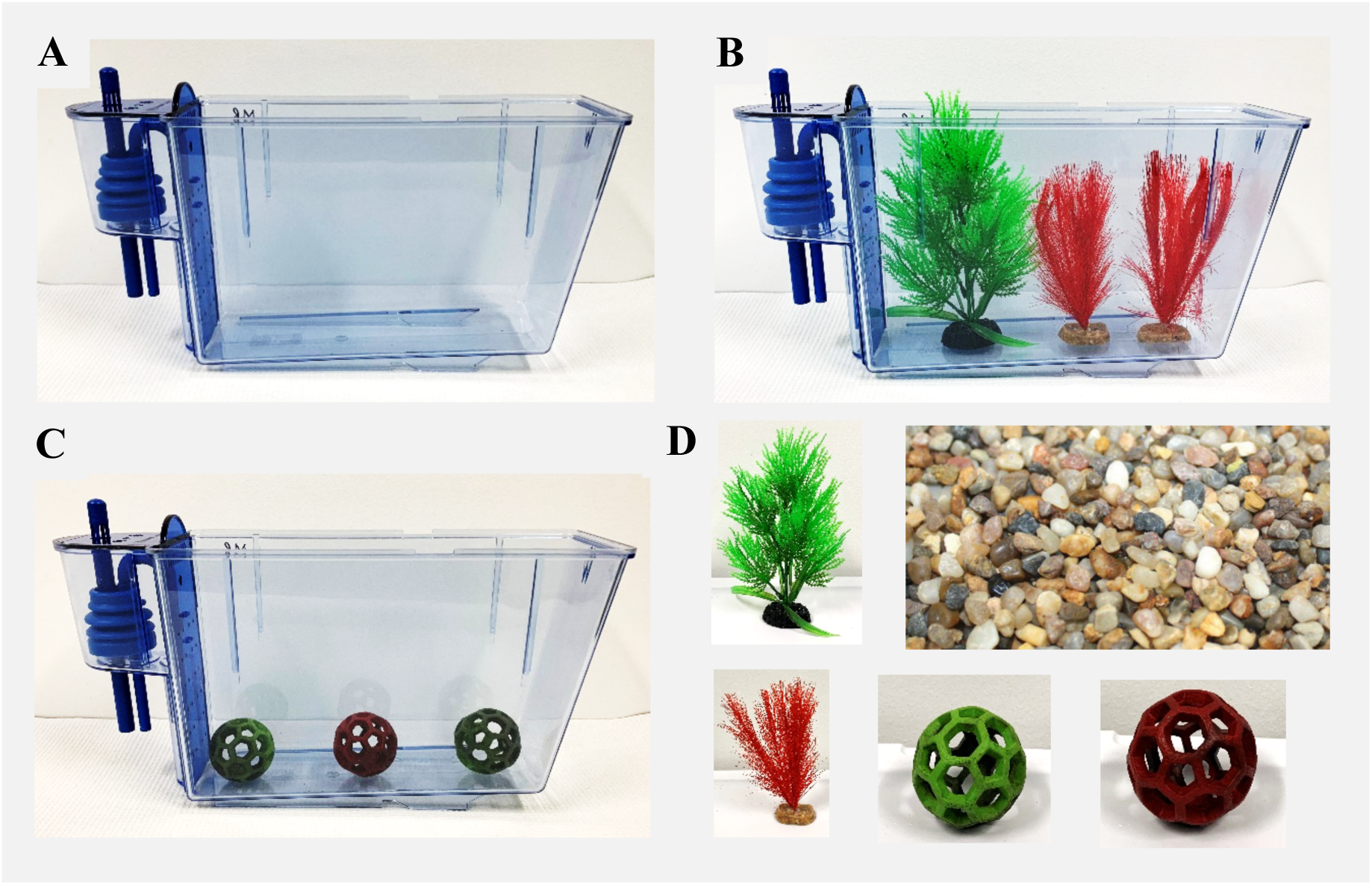
Experimental Tank Setup. **A.** The standard 3.5 L zebrafish housing tank which measures 27.2 cm x 10.2 cm x 17 cm. This type of tank housed all fish in the experiment. Fish from the control group and the gravel experimental group used an empty tank. **B.** Fish in the plant experimental group used a tank with artificial plants added. **C.** Fish in the toy experimental group used a tank with rubber toys added. **D.** The standalone enrichment items used in the experimental tanks. Each tank was separated by an opaque laminated screen. The gravel group(s) had the displayed image of gravel, measuring 20.8 cm x 26.8 cm, printed on their side of the screen. The other groups had a plain grey background on their side of the screen. The fish were housed on a single row of a Tecniplast ZebTec Active Blue system with an automatically circulating system.

#### Aggression Tests

In-tank aggression is difficult to monitor and record, however, a zebrafish’s level of aggression can be deduced using a mirror test (Woodward et al., 2019). The test consists of a single adult fish, the fish were 18 months old at this point, being held in an enclosure with a mirror at one end. We used a Tecniplast zebrafish 1.0 L breeding tank for the enclosure and placed it inside a matte white box to reduce distractions while filming. The trials were filmed with a Logitech C920x HD Pro Webcam at 30 frames per second. The fish was allowed to acclimate to the test tank for at least 10 minutes before the tank was moved to the white box serving as the experimental arena. They were then recorded from above for 10 minutes with the mirror revealed after 5 minutes had elapsed. Their movement was tracked from the acquired videos using DeepLabCut (Mathis et al., 2018) and their locations in coordinates for each frame were analyzed with a custom MATLAB script. The amount of time they spent in one of the three equally divided sections: closest to the mirror, furthest from the mirror, and in the center of the arena, was measured and used to determine their aggression (**Figure 3**). Increased proximity to the mirror after it is revealed is used as an indication of increased aggression (Woodward et al., 2019). Aggression can also be observed in attacks made by the fish toward its mirror image, however, this behavior does not always occur during these types of tests and was not quantified during the trials. Six fish were randomly selected from each tank and recorded individually for a single trial providing six total trials for each group.

**Figure 2.**
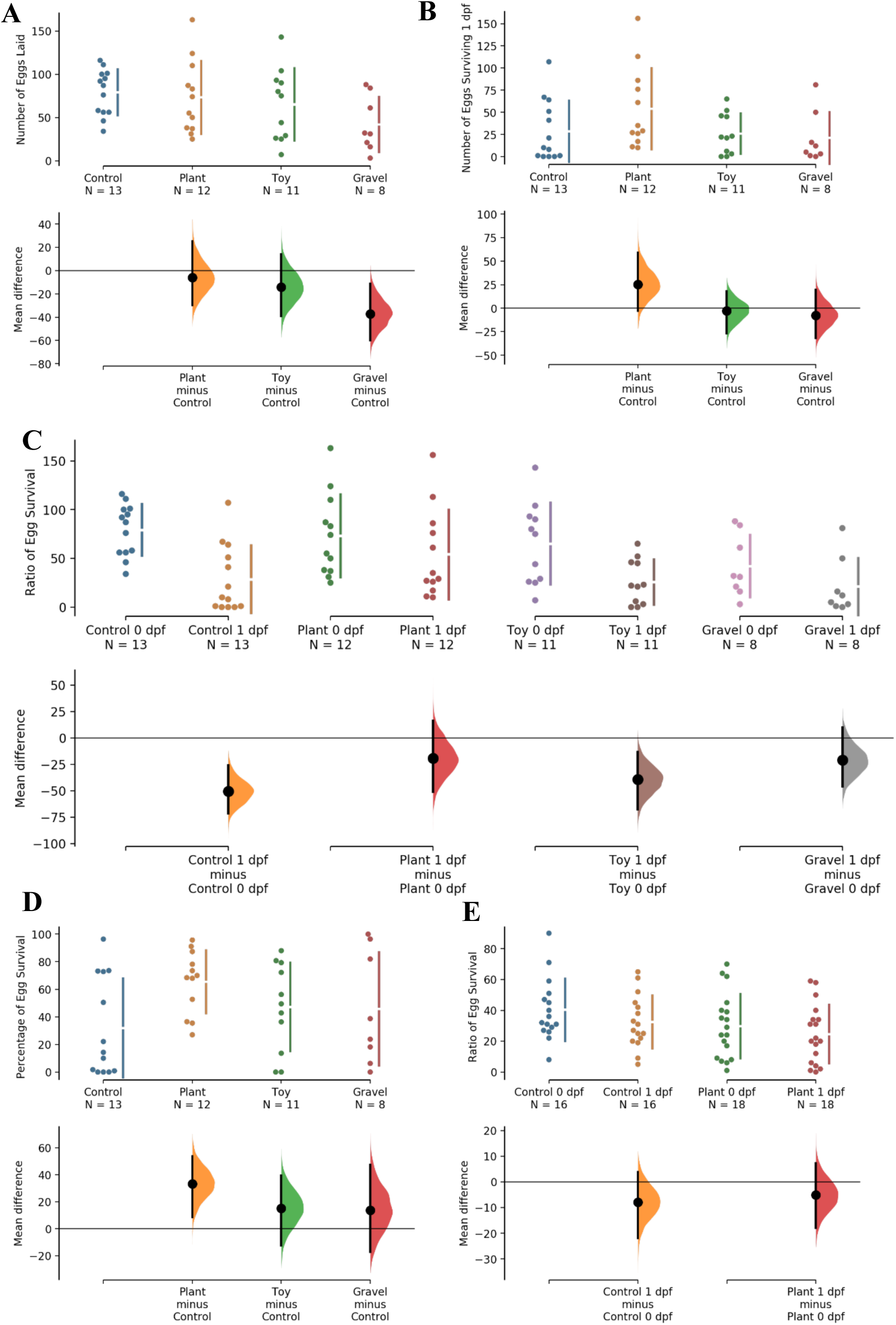
Effect of environmental enrichment on reproduction. Estimation statistics figures displaying the relevant results of the long-term and short-term exposure reproductive tests. In the upper part of the plot, each point represents a trial with N indicating the number of trials. The lower part of the plot uses bootstrap resampling to show the distribution of a larger sample. The middle line represents the independent variable that the bootstraps can be compared to. **A**, **B**, **D**, and **E** were generated using shared control. **C** was generated using multiple two-groups. **A**. Number of eggs laid. **B**. Number of eggs surviving 1 dpf. **C**. Ratio of egg survival. **D**. Percentage of egg survival. **E**. Ratio of egg survival from short term tests.

**Figure 3.**
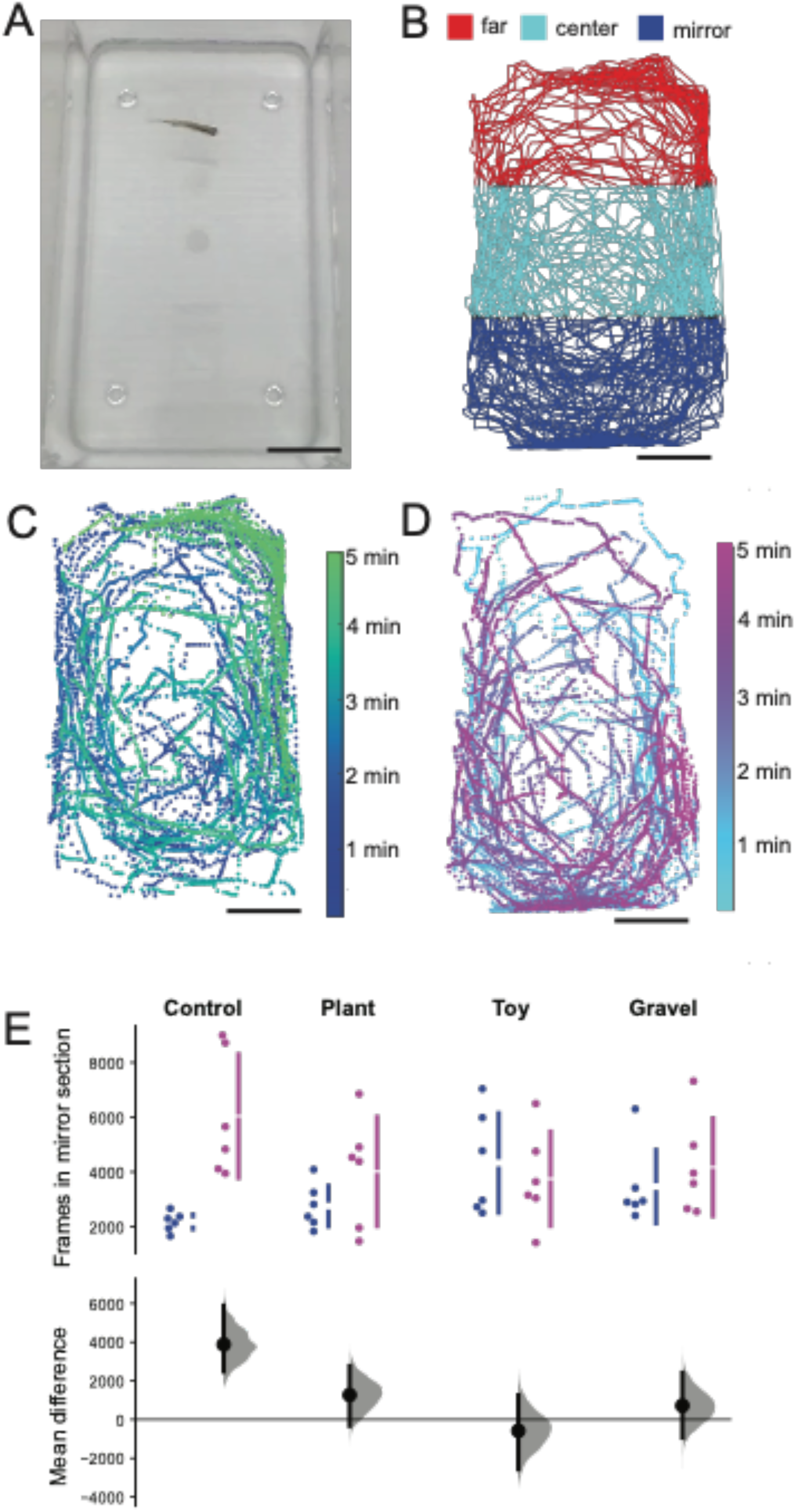
Effect of environmental enrichment on aggression. A. Single raw frame from a video recording of a fish in the arena. The mirror was revealed at the end of the tank shown at the bottom of the image. B. Raw traces of a tracked fish trajectory color coded by arena section depending on proximity to the mirror. The furthest third of the arena from the mirror is denoted as far (red), the center of the arena (cyan), and the third of the arena closest to the mirror side is denoted as mirror (blue). C. Position of a fish for each frame as the trial progresses, with data taken from the first half of the trial only, before the mirror is revealed. D. Position of a fish for each frame as the trial progresses, with data taken from the second half of the trial only, after the mirror is revealed. E. Estimation statistics comparing across treatment groups for the number of frames spend in the section closest to the mirror. (Top) Point in blue are the average number of frames per fish from before the mirror is revealed, where point in magenta are from after the mirror is revealed. N= 6 fish per treatment group. (Bottom) Mean difference between the data before and after the mirror is revealed for each treatment group. Scale bar in A-D 3 cm.

#### Breeding Tests

Fecundity or breeding effectiveness can be measured as success in laying eggs when the opportunity arises and success in producing eggs which can survive until the embryo hatches. We defined survivability as the number of eggs produced from which embryos survive until hatching divided by the total amount of eggs laid. We defined spawn rate as the number of times a pair produced eggs divided by the number of breeding encounters the pair is given. Overall breeding success was calculated using both factors (**Figure 2**).

The breeding tests consisted of a series of crosses where two fish were isolated in a Tecniplast USA 1.0 L breeding tank. Individuals from each group of fish were bred with other members of their group. The number of eggs collected from each cross were recorded as well as the number of eggs which survived to 1 dpf. 13 pairs from each group were bred twice, then measured and sexed again after their second cross (**Table 2**, **Table 3**).

**Table 1.** Fish Total Length Measurements (cm) and Sex Identification of all Experimental Fish at 100 dpf. All the fish which survived until 100 dpf were visually sexed and measured.

| **Group** | **Control** | **Plant** | **Toy** | **Gravel** |
|---|---|---|---|---|
| **Mean** | 2.49 | 2.49 | 2.44 | 2.37 |
| **Min** | 1.7 | 1.3 | 2 | 1.6 |
| **Max** | 2.9 | 2.8 | 2.7 | 2.9 |
| **Range** | 1.2 | 1.5 | 0.7 | 1.3 |
| **Mode** | 2.4 | 2.5 | 2.4 | 2.4 |
| **# of Females** | 25 | 17 | 20 | 21 |
| **# of Males** | 9 | 19 | 19 | 22 |

**Table 2.**
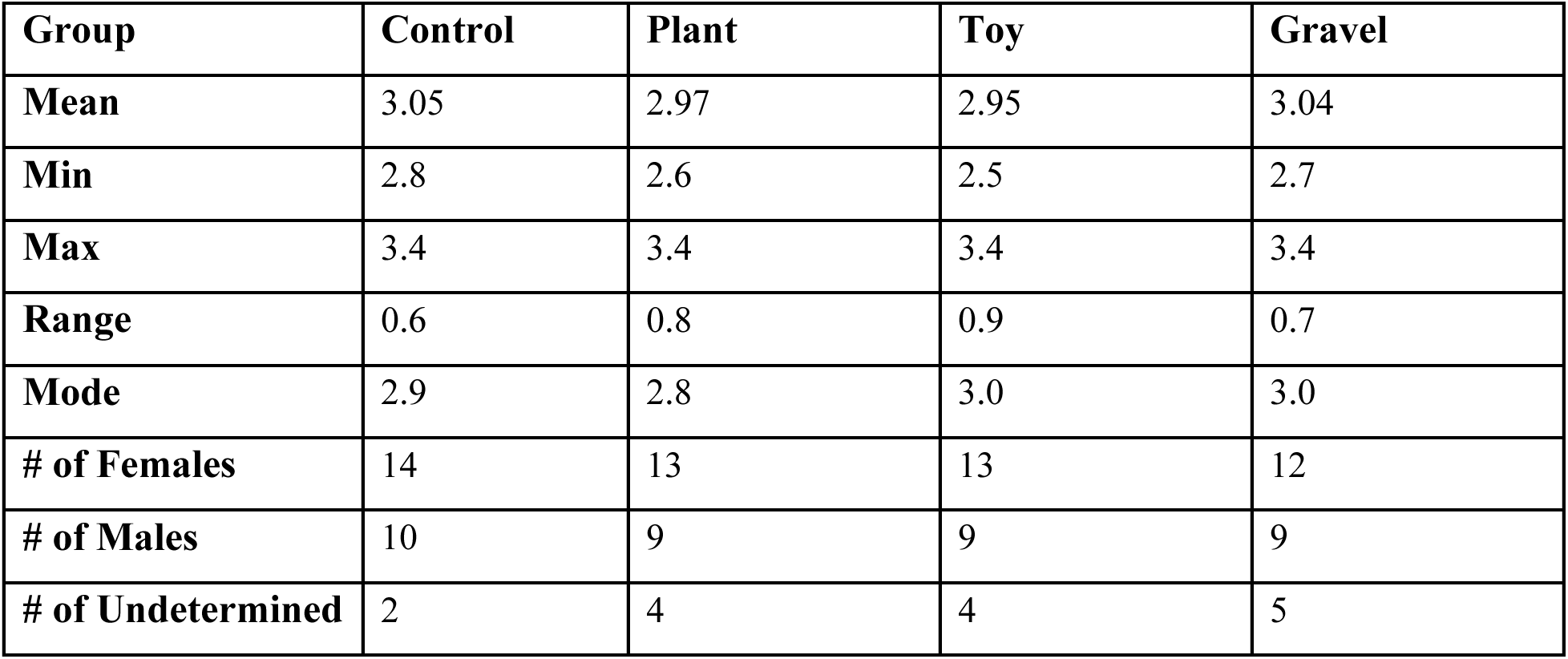
Fish Total Length Measurements (cm) and Sex Identification of Breeding Fish after 245 dpf. Measurements and sex identification after each group of fish completed their second attempt at spawning. The last three row are the quantity of female and male fish which could be identified, as well as the number of fish which could not be clearly identified as either male or female. 26 fish from each group were included in the breeding test and consequently measured and sexed. The last group of measurements were taken when the fish were at 266 dpf.

| **Group** | **Control** | **Plant** | **Toy** | **Gravel** |
|---|---|---|---|---|
| **Mean** | 3.05 | 2.97 | 2.95 | 3.04 |
| **Min** | 2.8 | 2.6 | 2.5 | 2.7 |
| **Max** | 3.4 | 3.4 | 3.4 | 3.4 |
| **Range** | 0.6 | 0.8 | 0.9 | 0.7 |
| **Mode** | 2.9 | 2.8 | 3.0 | 3.0 |
| **# of Females** | 14 | 13 | 13 | 12 |
| **# of Males** | 10 | 9 | 9 | 9 |
| **# of Undetermined** | 2 | 4 | 4 | 5 |

**Table 3.**
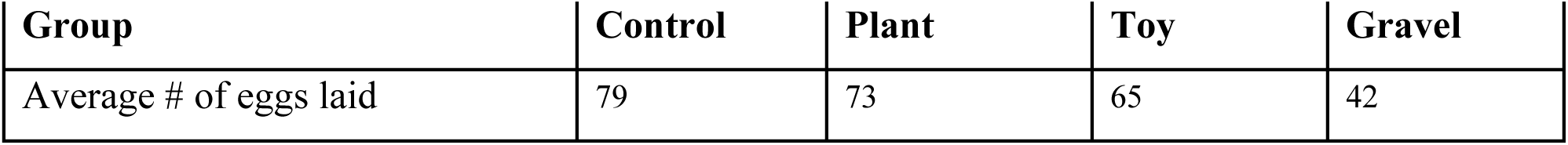
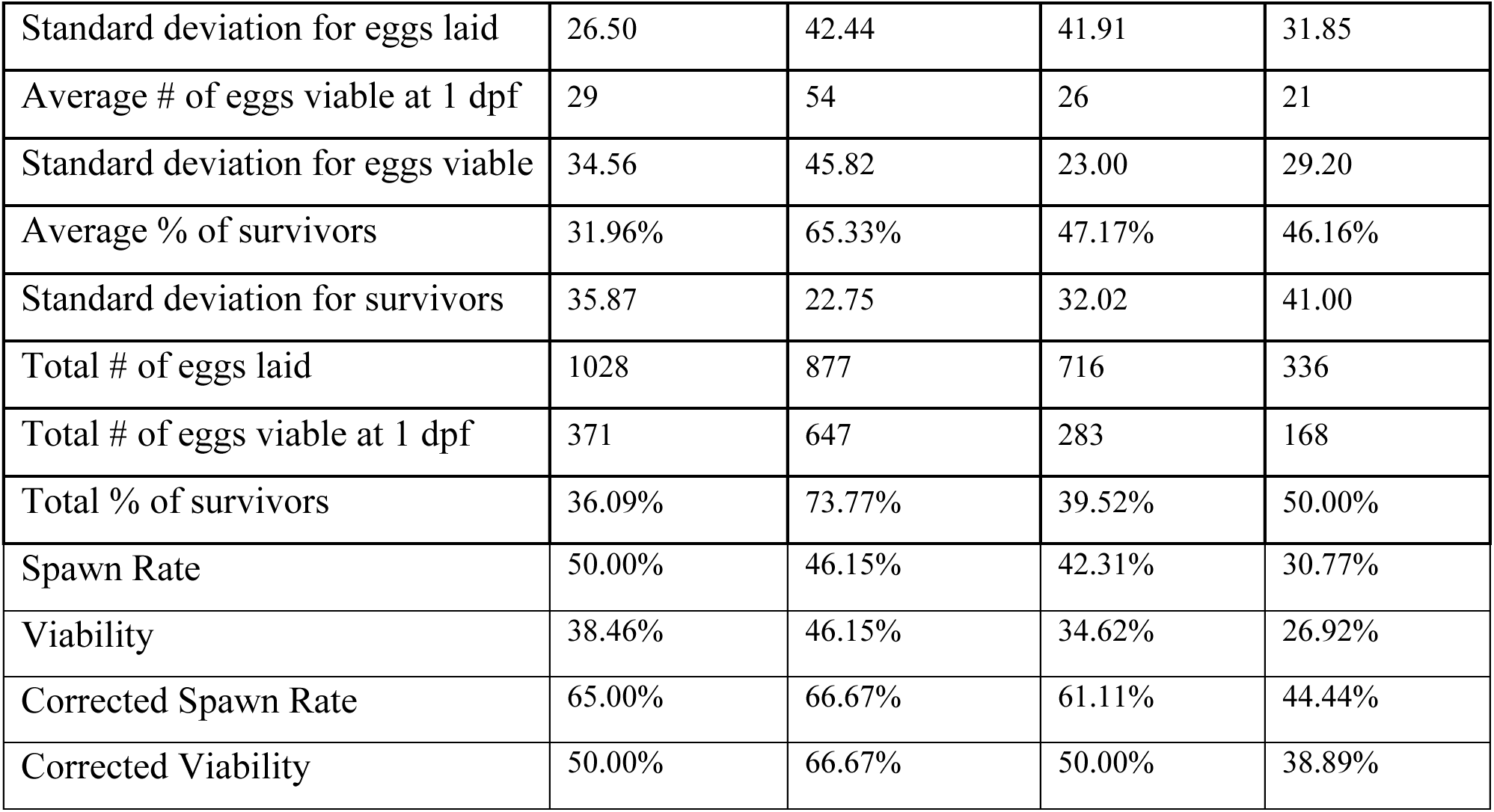
Effect of long-term environmental enrichment on breeding. The number next to the asterisk (*) denotes the standard deviation of the averages. The spawn rate is the percentage of crosses which resulted in eggs. The viability is the percentage of crosses which resulted in eggs that survived past 1 dpf. During the measurement and sexing after 245 dpf which directly followed the second round of tests, some of the fish could not be identified as either male or female with a high degree of certainty. For this reason, corrected values are also reporting the results from the reproduction tests only using data from the pairs that could be visually confirmed to contain one male and one female.

The ratio of eggs which survived until 1 dpf to eggs laid was defined as the survival. The ratio of breeding attempts which yielded eggs to the total number of breeding attempts made was defined as the spawn rate. A corrected spawn rate was calculated using the number of hypothetical breeding opportunities available to account for any human errors made when sexing the adult zebrafish. The number of male-female pairs which could be identified with a high degree of certainty served as the maximum number of hypothetical crosses. This number was used in place of the actual number of crosses for the “corrected” spawn rates. This information together created each groups’ breeding success rate (**Table 3**). The first round of breeding tests were conducted at 160 dpf and the second round at 244 dpf.

#### Measuring Growth

At 100 dpf the fish were anaesthetized with a solution of 168 mg/L Tricaine methane sulfonate (MS222, Tricaine-S, Thermo Fisher Scientific) and fish water from a stock solution made using 400 mg MS222 per 97.9 mL Reverse Osmosis, (RO), water and then buffered with 2.1 mL of 1M pH 9 Tris (Matthews and Vargas, 2012). The fish were then measured from nose to fin tip on a flat damp paper towel. The sex of the fish was also visually determined at this time. In cases where the sex could not be determined with a large degree of certainty, these were denoted separately (**Table 1**).

#### Short-term enrichment exposure Breeding Tests

After the conclusion of the aggression tests, additional breeding tests were conducted to assess if impact of the plant enrichment items on zebrafish egg survivability were the result of short-term or long-term exposure to the enrichment items. For these tests, fish which were raised in control conditions were bred in either control or enriched conditions before being returned to the control home tank. Male-female pairs of adult AB WT fish were randomly selected each week for the trials. The twelve pairs were then assigned to either a bare control breeding tank or an enrichment breeding tank with artificial plants affixed to the bottom. The artificial plants used in the breeding tanks were truncated versions of the plants used in the plant group housing tanks (**Figure 1**).

## Results

### Aggression Test Results

We used a mirror test to measure the amount of aggression in the adult zebrafish. The test presents the subject with a covered mirror at one end of their enclosure. Halfway through the trial, the mirror was revealed, and a significant increase in the amount of time spent in the area near the mirror was used as a mark of aggressive behavior. Statistical significance was established using an ANOVA test through estimation statistics. Long-term exposure to environmental enrichment significantly decreased aggressive behavior in zebrafish (**Figure 3**) as measured by the mirror test. Fish from the control group spent significantly more time near the mirror after it was revealed than the other three enrichment groups. Fish from the three enrichment groups had no significant difference in the amount of time spent in any of the three sections after the mirror’s reveal.

### Breeding Test Results

Thirteen pairs of fish from each experimental group were crossed twice to collect data for the breeding test. The number of eggs laid on the day of fertilization were recorded as well at the number of fertilized eggs which survived 1 day post fertilization. Statistical significance was established using an ANOVA test through estimation statistics. Long-term exposure to environmental enrichment significantly impacted the reproductive behavior of the zebrafish (**Table 3**, **Figure 2**), although not uniformly. Fish from the control group produced significantly more eggs overall than fish from the gravel group (p=0.0074, **Figure 2A**). Control fish egg production was not significantly higher than egg production of fish from the plant or toy groups. Control fish and gravel group fish also experienced a significant amount of offspring death between 0 dpf and 1 dpf which was not observed in the plant or toy group fish (p=0.0008 and p=0.0104 respectively, **Figure 2B**, **Figure 2C**). Fish from the plant group experienced significantly higher survival of offspring from 0 dpf to 1 dpf than fish from the control group, (p=0.0114), though it was not significantly higher survival than fish from the toy and gravel group (p=0.0114, **Figure 2D**). While several other trends can be observed from the data, these results are not statistically significant.

### Growth Test Results

We compared the sizes of the fish, measured in cm, from each group. They were anesthetized then measured on a damp surface using a standard ruler. Long-term exposure to environmental enrichment had no significant impact on the survivorship or growth of any of the experimental groups as measured at 100 dpf (Control vs plant p=0.970; Control vs toy p=0.294; Control vs gravel p=0.055) or after 250 dpf (Control vs plant p=0.097; Control vs toy p=0.054; Control vs gravel p=0.814). Though the ranges of the sizes of the fish in each group varied (**Table 1**, **Table 2**), there were no significant differences in the mean, median, mode or range of those values.

### Short-term Exposure Breeding Tests

Adult AB WT zebrafish raised separately under control conditions were bred with and without the presence of plant enrichment during breeding to assess the impact of short-term enrichment exposure on zebrafish reproduction. Statistical significance was tested using a *t*-test through the program estimation statistics. We observed no significant difference in the number of eggs laid or the number of eggs which survived until 1 dpf (**Table 4**, **Figure 2E**).

**Table 4.** Effect of short-term exposure to environmental enrichment on breeding. STD denotes the standard deviation of the averages. The spawn rate is the percentage of crosses which resulted in eggs. The viability is the percentage of crosses which resulted in eggs that survived past 1 dpf. The utility is the percentage of crosses which resulted in a number of ≥ n eggs that survived past 1 dpf.

| **Group** | **Control** | **Plant** |
|---|---|---|
| Average # of eggs laid | 40 (STD 19.51) | 29 (STD 20.33) |
| Average # of eggs viable at 1 dpf | 32 (STD 16.64) | 24 (STD 18.46) |
| Average % of survivors | 78.39% (STD 18.93%) | 75.03% (STD 25.30%) |
| Total # of eggs laid | 645 | 534 |
| Total # of eggs viable at 1 dpf | 519 | 442 |
| Total % of survivors | 80.47% | 82.77% |
| Spawn Rate | 66.67% | 75.00% |
| Viability | 66.67% | 70.83% |

## Discussion

These experiments specifically tested the long-term effects of enrichment on zebrafish behavior, but the effects of the enrichment items differed from what we hypothesized. We hypothesized long-term exposure to any of the three enrichment items would increase reproductive behavior in zebrafish, while also increasing aggressive behavior. Instead, we found highly variable impacts on reproductive measures and a significant decrease in aggressive behavior after rearing in enriched environments. Long-term exposure to the plant enrichment items increased the reproductive effectiveness of the zebrafish in this study in one respect, in that fish from the plant group experienced significantly higher survival of offspring than fish from the control group. During short-term exposure to plants during breeding had no significant impact. Perhaps most importantly, long-term exposure to all three types of enrichment items significantly reduced the amount of aggression in zebrafish as measured by a mirror test, which we would consider a potential enhancement to quality of life.

Since zebrafish are social and prefer living in communal groups, researchers often do not include enrichment in their tanks, as the presence of conspecifics themselves are considered enriching (Collymore et al., 2015). Beyond social housing, additional environmental enrichment is not always required by most regulatory bodies for housing zebrafish, but it can particularly improve the quality of life for a singly housed fish (Collymore et al., 2015). Zebrafish tanks are often kept bare to simplify laboratory routines such as cleaning the tanks, monitoring the health of the colony, and removing fish for breeding or experiments (Williams). While zebrafish can show a preference for enrichment, they prefer the company of their conspecifics (Collymore et al., 2015). This preference and the variable effects from the implementation of enrichment can make it difficult to argue for its presence to be mandatory in laboratory setups.

The experimental setup for the enrichment tanks posed several problems throughout the study. The additional items in the tank lowered visibility and made removal of fish from the enclosure difficult. The enrichment items also appeared to increase accumulation of debris inside the tank and while this variable was never quantified during the experiment, it was frequently noted by other researchers on the team (B. Rejouis pers. obs.). The different enrichment items also had heterogeneous effects on the fish, which made evaluating the overall usefulness of enrichment difficult. Most notably the experimental group which used gravel wallpaper as an enrichment item significantly underperformed in egg production compared to the control. Meanwhile the experimental group which used artificial plants outperformed the control when it came to egg survivability. While the breeding test yielded variable results, the mirror tests revealed that long-term exposure to enrichment items of any type significantly decreased the level of aggression in our study toward conspecifics.

Similar studies on enrichment’s effect on reproduction found results somewhat different than those observed in this experiment. Studies on the effect of short-term exposure on zebrafish reproduction found that the presence of plant type enrichment items significantly increased the number of eggs produced, however, these results were not consistent across all types of plant enrichment (Wafer et al., 2016; Woodward et al., 2019). Another study did not find any significant impact of plant type enrichment on zebrafish reproduction (Lee et al., 2018). This suggests the type of plant used for enrichment has as much importance as its presence when assessing its impact on zebrafish reproduction. However, these differences in results could also be caused by differences in the basic husbandry protocols of the research facilities in each study.

The goal of this study was to assess the benefit of implementing environmental enrichment items in zebrafish enclosures. Zebrafish husbandry has been long established and changes to the standard must come with significant benefits. Increased fecundity would reduce the amount of research animals needed overall, and decreased aggression would lower the risk of the subjects experiencing unnecessary harm. However, there are other factors that influence zebrafish behavior which are already considered essential to zebrafish husbandry. A zebrafish’s diet has significant effects on its fertility and fecundity and this is well documented (Fowler et al., 2019; Kolb et al., 2018; Lawrence et al., 2012; Monteiro et al., 2018). There are also numerous studies which explored the effects of environmental factors other than enrichment items have on zebrafish reproductive behavior such as stocking density, water quality, and light cycles (Adatto et al., 2016; Delomas and Dabrowski, 2018; Kolb et al., 2018; Martinez-Sales et al., 2014; Peng et al., 2019). These factors are integral to zebrafish husbandry, yet their parameters vary between different facilities (Kolb et al., 2018).

We believe there is a greater need for standardization of husbandry methods for laboratory zebrafish. The knowledge gained from research on these animals should also be reflected in the methods used to house them. The results of this study reject the notion that bare tanks are the optimal housing environment for laboratory zebrafish. Finding the ideal conditions to raise zebrafish in would not only improve their quality of life, but also improve the quality of research results by encouraging the animals to mimic the behaviors observed when not in captivity. Several facilities have already started studying how varying amount of food, space, and heat can affect zebrafish behavior (Adatto et al., 2016; Delomas and Dabrowski, 2018; Fowler et al., 2019; Kolb et al., 2018; Lawrence et al., 2012; Monteiro et al., 2018). Future efforts should also focus on identifying which categories of environmental enrichment elicit beneficial actions beyond those used in this experiment and finding the most effective method of implementing those aspects.

## Acknowledgements

We would like to thank and acknowledge the following for their contributions to this research project: Dr. Shivon Boodhoo for useful discussion and insisting my own questions about zebrafish enrichment should be personally explored and answered. Dr. Caroline DeVan for feedback and guidance on the experimental methods and the parameters that should be tested. The Fortune Lab team for use of their Deep Lab Cut setup. Dr. Eric Fortune for advice the IACUC proposals and amendments needed to conduct the experiment, analysis of the mirror test videos using Deep Lab Cut, and MATLAB code needed to interpret the results of the mirror test videos. Finally, the Severi Lab team for assistance with fish care and feeding throughout the project and breeding of the zebrafish observed during the project.

## Author Contributions

BCR conceived of the project with guidance from KES. BCR performed all experiments and measurements. BCR and KES analyzed the data, made the figures, and wrote the manuscript.

## Protocol registration

No protocol was registered.

## Data access

All data and code are available upon request.

## Abbreviations and Acronyms

Dpf: days post fertilization

